# The Cx43 Carboxyl-Terminal Mimetic Peptide αCT1 Protects Endothelial Barrier Function in a ZO1 Binding-Competent Manner

**DOI:** 10.1101/2021.06.19.449103

**Authors:** Randy E. Strauss, Louisa Mezache, Rengasayee Veeraraghavan, Robert G. Gourdie

## Abstract

The Cx43 CT mimetic peptide, αCT1, originally designed to bind to ZO1 and thereby inhibit Cx43/ZO1 interaction, was used as a tool to probe the role of Cx43/ZO1 association in regulation of epithelial/endothelial barrier function. Using both *in vitro* and *ex vivo* methods of barrier function measurement, including Electric Cell-Substrate Impedance Sensing(ECIS), a FITC-dextran transwell permeability assay, and a FITC-dextran cardiovascular leakage protocol involving Langendorff-perfused mouse hearts, αCT1 was found to protect the endothelium from thrombin-induced breakdown in cell-cell contacts. Barrier protection was accompanied by significant remodeling of the F-actin cytoskeleton, characterized by a redistribution of F-actin away from the cytoplasmic and nuclear regions of the cell, towards the endothelial cell periphery, in association with alterations in cellular orientation distribution. In line with observations of increased cortical F-actin, αCT1 upregulated cell-cell border localization of endothelial VE-cadherin, the Tight Junction protein Zonula Occludens 1 (ZO1), and the Gap Junction Protein (GJ) Connexin43 (Cx43). A ZO1-binding-incompetent variant of αCT1, αCT1-I, indicated that these effects on barrier function and barrier-associated proteins, were likely associated with Cx43 CT sequences retaining ability to interact with ZO1. These results implicate the Cx43 CT and its interaction with ZO1, in the regulation of endothelial barrier function, while revealing the therapeutic potential of αCT1 in the treatment of vascular edema.

## Introduction

Barrier function is a vital mechanism characterized by the homeostatic exchange of substances between interior and exterior compartments of epithelial tissues, marked by apical and basolateral membrane domains, respectively (Zihni, Mills, Matter, & Balda, 2016). Diseases associated with vascular barrier function disruption occur in the heart and other tissues, the functions of which critically depend upon a healthy blood circulation. These diseases include ischemia-reperfusion Injury, coronary artery disease (CAD), stroke, acute respiratory distress syndrome (ARDS), chronic skin wounds such diabetic foot ulcers as well as many other pathologies (Aghajanian et al., 2008; Escribano et al., 2019; Herrero, Sanchez, & Lorente, 2017; Gerd Heusch, 2016; Heusch, 2018; Higashi & Miller, 2017; Simmons, Erfinanda, Bartz, & Kuebler, 2019; Soon, Chua, & Becker, 2016). The vascular endothelial barrier, a specialized epithelial monolayer lining blood vessels, acts like a semi-permeable filter that regulates the exchange of cells, extracellular vesicles, plasma proteins, solutes, and fluids between the circulation and tissue (Aghajanian, Wittchen, Allingham, Garrett, & Burridge, 2008; Komarova & Malik, 2010). Pathological stress triggers breakdown in these barrier properties, causing characteristic disruptions in cytoskeletal structure and junctional complexes at cell-cell contacts, including intercellular gap formation (Belvitch, Htwe, Brown, & Dudek, 2018). These changes can result in edematous buildup of fluid, ions and other solutes, as well as enhanced immune cell infiltration across multiple tissue types and disease processes (Aghajanian et al., 2008; Escribano et al., 2019).

Cellular structures involved in regulating barrier function include: 1) The Tight junction (TJ), which provides a gating mechanism that directly controls the exchange of substances across the paracellular space; 2) The Adherens junction (AJ), which is critical for the establishment and maintenance of cell-cell adhesion; 3) The Actin cytoskeleton, which controls the overall integrity of cell-cell contacts via mechanical push*/*pull forces; and (4) The Gap junction (GJ), which allows for exchange of signaling molecules and ions between cells through connexin-based transcellular channels, in addition to providing close points of intercellular adhesion (Derangeon, Spray, Bourmeyster, Sarrouilhe, & Hervé, 2009; B. N. Giepmans, 2004; Radeva & Waschke, 2018). While initially conceived of as independent, these transcellular complexes were subsequently identified as sharing direct interactions with the tight junction scaffolding molecule, Zonula Occludens 1 (ZO1), which is thought to contribute to biochemical and biophysical crosstalk between their protein components (Derangeon et al., 2009; Garcia, 2009; B. N. Giepmans, 2004; Hervé, Bourmeyster, Sarrouilhe, & Duffy, 2007).

Findings have emerged over the last 20 years or more that gap junctional connexins, especially the most studied isoform, Connexin 43 (Cx43), influences barrier function and permeability (Strauss & Gourdie, 2020). There is also growing appreciation that this may involve both channel-dependent and independent functions of Cx43, including via effects on intercellular communication, membrane permeability, cell-cell contact arrangements and cytoskeletal dynamics, junction assembly, cell polarity, and transcriptional regulation (Francis et al., 2011; Kameritsch, Pogoda, & Pohl, 2012; Leithe, Mesnil, & Aasen, 2018; Matsuuchi & Naus, 2013; Olk, Zoidl, & Dermietzel, 2009). While mounting evidence suggests that Cx43-based channel activity can modulate barrier function changes under pathological stress conditions, the channel-independent role of Cx43 in barrier modulation is less understood. The Cx43 carboxyl-terminus (CT), exhibits a well-characterized interaction with ZO1, specifically at its PDZ2 domain (B. N. G. Giepmans & Moolenaar, 1998; Sorgen et al., 2004; Sorgen, Trease, Spagnol, Delmar, & Nielsen, 2018). While the details of this structural interaction are well-established, the functional consequences remain to be characterized. In this study, we examine the effects of short mimetic peptides based on the Cx43 CT sequence, with and without the capacity to interact with ZO1. Our results indicate that αCT1, which incorporates the CT-most 9 amino acids of Cx43, protects endothelial cell barrier function in a ZO1 interaction-associated manner. αCT1 is presently in clinical testing in humans for healing of normal and chronic skin wounds (Lampe and Laird, 2018; Strauss and Gourdie, 2020). The barrier protective effect of αCT1 is accompanied by marked changes in patterns of ZO1, VE-Cadherin, Cx43, and actin cytoskeleton remodeling in peptide-treated cells. Taken together, our data suggests that modulation of actin-based inter- and intra-cellular push/pull forces may be a key aspect of the molecular mechanism of αCT1 on barrier function, contributing to the mode-of-action of this therapeutic peptide in regulating tissue edema.

## Results

### αCT1 requires a CT isoleucine to associate with ZO1 at borders between MDCK cells

The αCT1 peptide consists of the CT-most 9 amino acids of Cx43: Arg-Pro-Arg-Pro-Asp-Asp-Leu-Glu-Iso or RPRPDDLEI, includes a 16–amino acid N-terminal antennapedia sequence (ANT) and typically has an N-terminal biotin tag (Hunter, Barker, Zhu, & Gourdie, 2005). The last four amino acids of αCT1 (DLEI) mimic the class II PDZ-binding motif of Cx43, which has been shown to mediate interactions with the second of the three PDZ (PDZ2) domains of the tight junction protein, ZO1 (Jiang et al., 2019). Deletion of the CT isoleucine of this motif abrogates interaction of αCT1 with ZO1 PDZ2, as is the case with the ZO1 binding-incompetent αCT1-I variant used in this, as well as our previous work on the molecular mechanism Cx43 CT peptides in mitigating cardiac ischemia reperfusion injury (Jiang et al., 2019). In these studies, we demonstrated that in addition to interacting with ZO1, αCT1 has the capacity to interact with Cx43 CT itself.

Using Electric Cell-Substrate Impedance Sensing (ECIS) we previously reported that αCT1 abrogates EGTA-induced loss of barrier function in retinal pigment epithelial monolayers (Obert et al., 2017). Follow-up ECIS experiments in the present study indicated that Cx43-deficient Madin-Darby Canine Kidney (MDCK) cell cultures were similarly protected by αCT1, but not αCT1-I, from a Ca^2+^ chelating, EGTA-treatment. The addition of 100µM αCT1, 5 min after Ca2+ chelation with 2mM EGTA, produced barrier function recovery beyond that observed with the control peptides, αCT1-I, and the cell penetration sequence alone, antennapedia (ANT) (Supplementary Figure 1).

These initial barrier function findings in MDCK cells demonstrated the barrier function-modulating potential of the 9 amino acid (aa) CT-most sequence of Cx43. These observations further indicated that αCT1’s mechanism of action likely involved ZO1 binding-competency. To confirm that αCT1 interacts with ZO1 inside the cell, αCT1’s association with the tight junction protein, ZO1, was investigated. We first examined αCT1 uptake and distribution in MDCK cells using confocal microscopy. Cx43-negative MDCK cells (Figure 1A) were used in this analysis to reduce confounding binding of the αCT1 and αCT1-I to Cx43 itself, a characteristic of both peptides that we have demonstrated previously (Jiang et al., 2019). Consistent with results from HeLa cells (Hunter et al., 2005), we observed robust antennapedia peptide-mediated uptake into MDCK cells. However, unlike HeLa cells, MDCK cells incubated with αCT1 showed dense concentrations of peptide co-localized with ZO1 at cell-cell borders (Figure 1C). This pattern appeared to occur in a dose-responsive manner (Figure 1B), with signal intensity increasing with increasing concentrations of applied αCT1 - from 5µM, 100µM to 150µM. This distinctive co-localization is illustrated further in a 3D-volumetric rendering in Figure 2B, where αCT1, but not αCT1-I, can be seen to uniformly and intensely co-localize with ZO1 at an interface containing the tight junction belt between apposed cells. Quantitative analyses confirmed that αCT1 colocalized with ZO1 at cell borders at significantly higher levels than the ZO1 binding-incompetent peptide αCT1-I, antennapedia (ANT) peptide alone (i.e., with no additional Cx43-related sequence), and vehicle controls, with no added peptide (Fig 2A, C). To further substantiate αCT1-ZO1 association, we performed proximity ligation assays (Duolink) using antibodies against ZO1 and the biotin tags present on αCT1 and αCT1-I peptides. Punctate ZO1-biotin Duolink signals were significantly increased following incubation of cells with αCT1, but largely attenuated following incubation with αCT1-I (Figure 2D, E). Taken together, these data suggested that αCT1 associates in close proximity with ZO1 at Cx43-negative MDCK cell-cell borders, requiring a functional ZO1 PDZ2 domain-binding motif to maintain this pattern. These results were consistent αCT1 directly targeting and binding to ZO1 located at cell-to-cell junctions.

**Figure 1:**
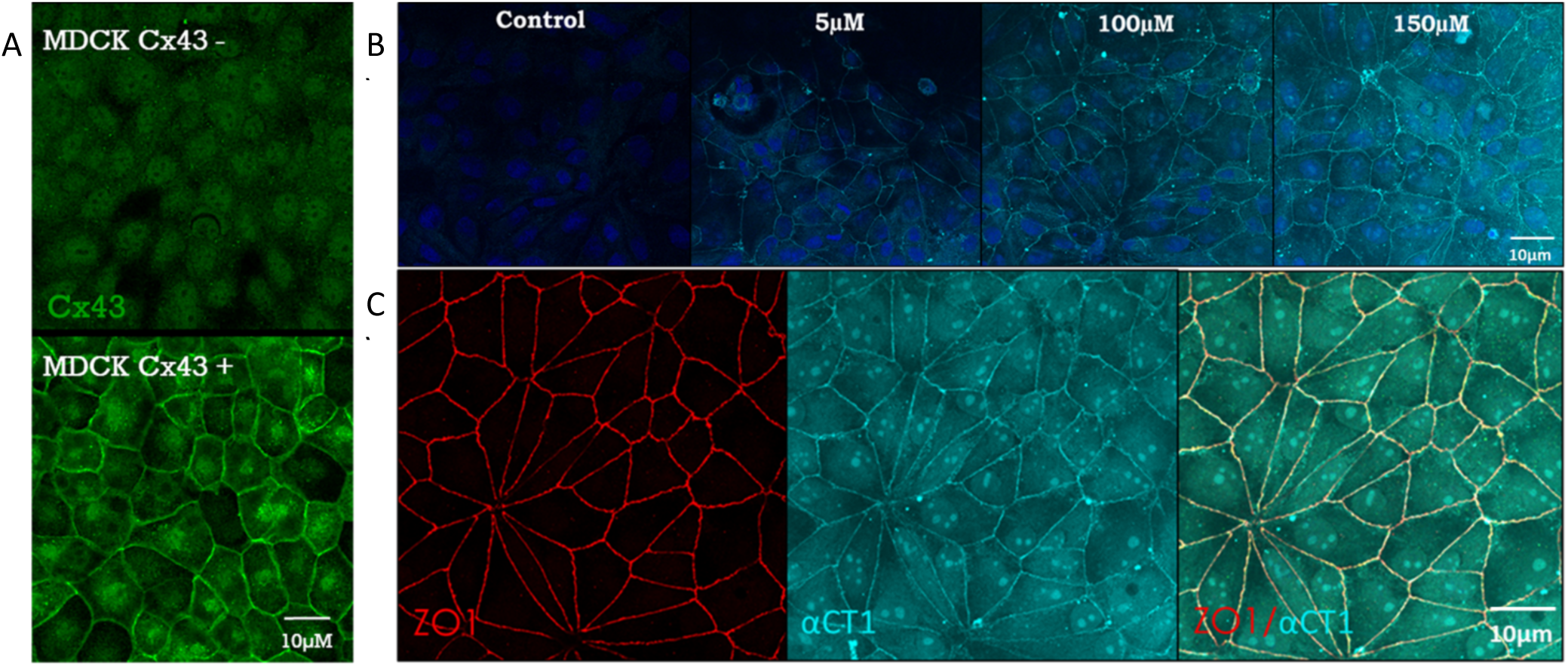
Cx43 CT mimetic peptide, αCT1, colocalizes with ZO1 inside Cx43-deficient MDCK cells. **A)** Representative confocal images of Cx43-deficient and Cx43-expressing MDCK cells. **B)** Representative confocal images of the dose-dependent uptake of fluorescent streptavidin-labeled, biotinylated αCT1 (0, 5, 100,150 µM) inside Cx43-deficient MDCK cells, fixed at 1 h post-incubation of peptide. **C)** Representative confocal images of ZO1 + αCT1, combined into a merged image to highlight colocalization (yellow).

**Figure 2:**
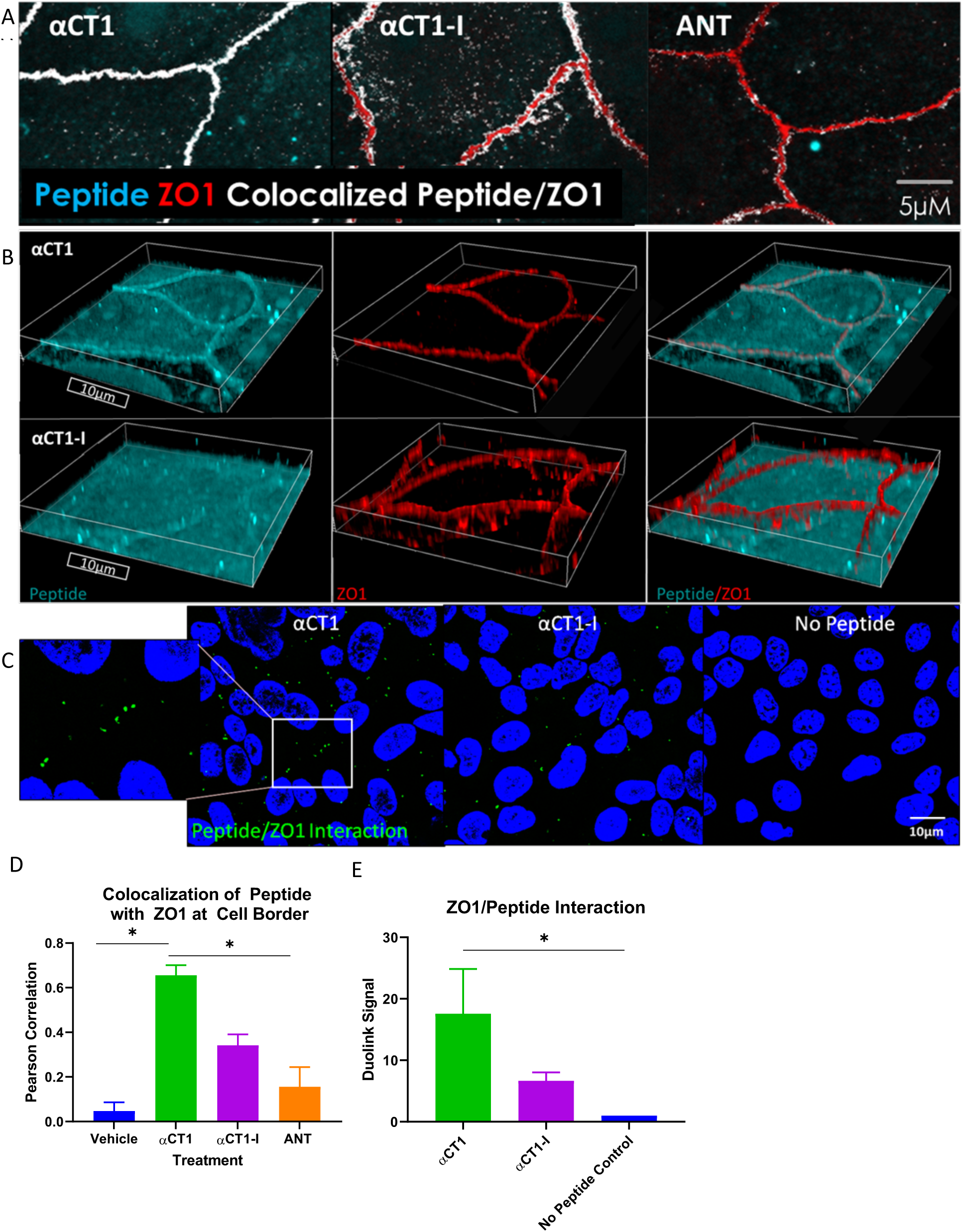
αCT1 requires its terminal isoleucine to associate with ZO1 inside Cx43-deficient MDCK cells. **A)** Representative confocal images of colocalization (white) between αCT1 and ZO1 binding-deficient control, αCT1-I, and cell penetration sequence control, antennapedia (ANT). **B)** Representative volumetric 3D confocal image renderings of border localization of αCT1 vs αCT1-I. **C)** Representative confocal images of the Duolink interaction between peptides and ZO1. Green spots represent points of interaction. **D)** Quantification of colocalization between the peptides and ZO1, as determined by Pearson Correlation analysis. **E)** Quantification of the Duolink interaction between the peptides and ZO1. *P < 0.05 vs. controls; N = 3.

### αCT1 inhibits thrombin-induced disruption of endothelial barrier function in a ZO1 binding-competent manner

Previous reports have demonstrated that the Cx43 mimetic, αCT1, has cardioprotective properties in an *ex-vivo* mouse model of global ischemia reperfusion injury (Jiang et al., 2019). We considered that targeting of the coronary vasculature and effects on edema were an unexplored aspect of the mode-of-action αCT1 in cardioprotection. Breakdown in endothelial barrier function is a hallmark of several cardiac pathologies, including ischemia-reperfusion injury (Heusch, 2018; Mezache et al., 2020). The barrier protective effects observed in the MDCK cells (Supplementary Figure 1) raised the possibility that αCT1 might similarly protect barrier function within endothelial cells. To investigate the potential for αCT1 to protect endothelial barrier function, we used ECIS to assess the barrier-modulating effect of αCT1 and the ZO1-binding incompetent control αCT1-I in microvascular endothelial cell monolayers. To this end, confluent HMEC-1 monolayers were grown on ECIS electrode arrays, treated with peptide (100µM) for 1h, then stimulated with thrombin (1U/mL). Thrombin is a well-known barrier function disruptor and bona fide inflammatory mediator of ischemia reperfusion injury and other cardiac diseases (Jackson, Darbousset, & Schoenwaelder, 2019). ECIS indicated that pretreatment with αCT1, but not αCT1-I, significantly attenuated barrier function disruption induced by thrombin in HMEC-1 monolayers (Figure 3A, B). Interestingly, we noted from ECIS records that a significant level of stabilization occurred prior to treatment with thrombin, during the one hour period in which cells were incubated with αCT1 (Figure 3C). Again, a similar pre-treatment effect was not observed for αCT1-I (Figure 3C). To further validate our results, we repeated the experiment using a second well-characterized assay of barrier integrity, a transwell permeability assay based on the flux of a 4.5 kDa TRITC dextran permeability tracer across the monolayer (Figure 4). In line with the ECIS data, αCT1 pretreatment significantly blocked hyperpermeability to the tracer following exposure to thrombin – at the 10 minute time point of maximum disruption (Figure 4B), as indicated from initial time course experiments (Figure 4A). By contrast, αCT1-I demonstrated no barrier protecting effect. Based on these data we concluded that pretreatment with αCT1 was sufficient to maintain endothelial barrier in the context of thrombin-induced disruption, whereas the ZO1-binding incompetent variant peptide αCT1-I was unable to mediate a similar protective effect.

**Figure 3:**
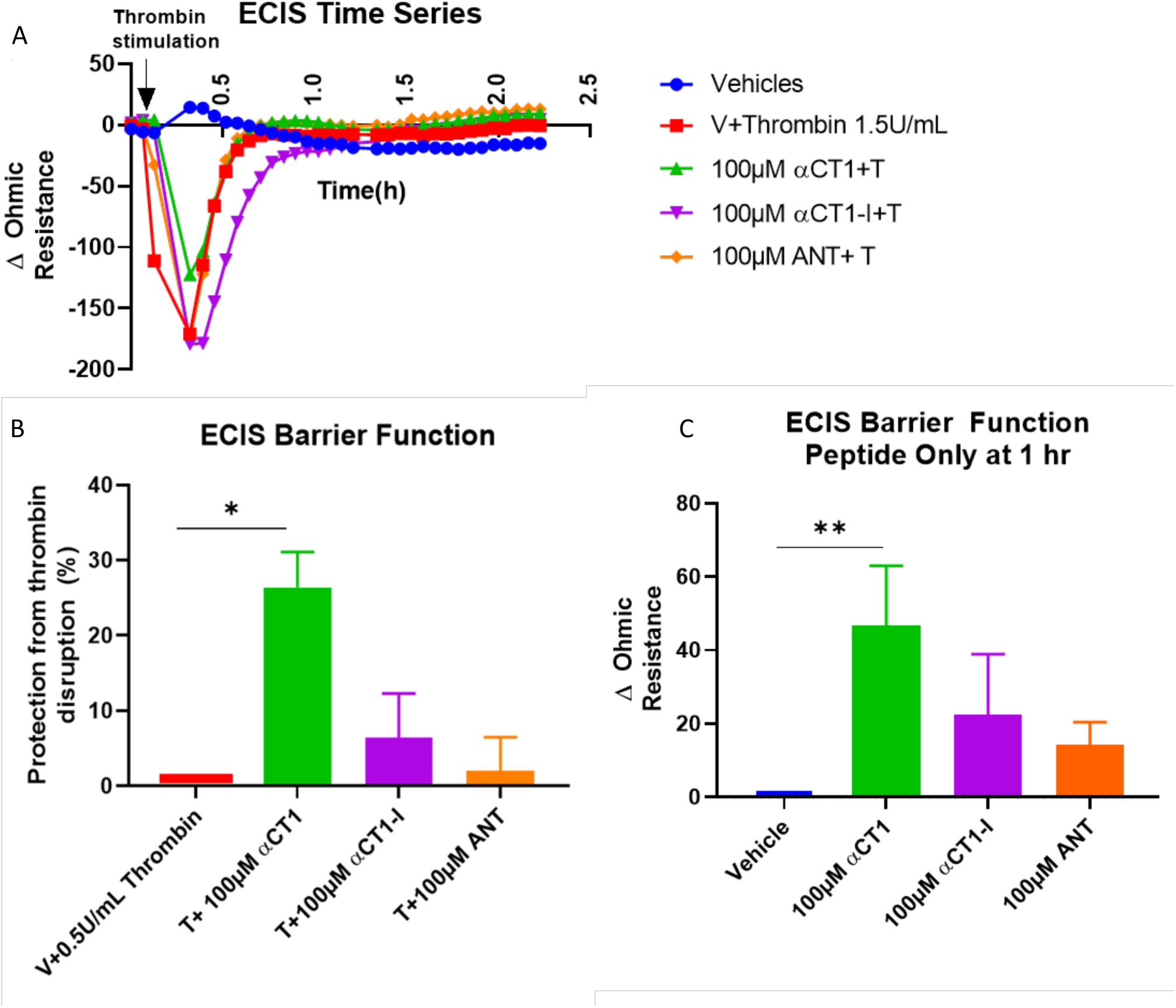
αCT1 requires ZO1 binding-competency to protect the endothelial barrier from thrombin-induced disruption measured by Electric Cell-substrate Impedance Sensing (ECIS) in HMEC-1 cell monolayers. **A)** Representative ECIS time series showing peptide-induced barrier function changes following thrombin treatment. Each data point represents the change in ohmic resistance from individual treatment baselines, collected at approx. 4 min intervals. **B)** Approximately 5 min following thrombin addition, peptide-induced barrier function protection was calculated as the percentage of barrier protection from thrombin disruption **C)** At 1 h peptide incubation, prior to thrombin treatment, the effects of peptides on barrier function were calculated as the change in ohmic resistance compared to vehicle control. * P < 0.05 vs. controls; N = 3-5.

**Figure 4:**
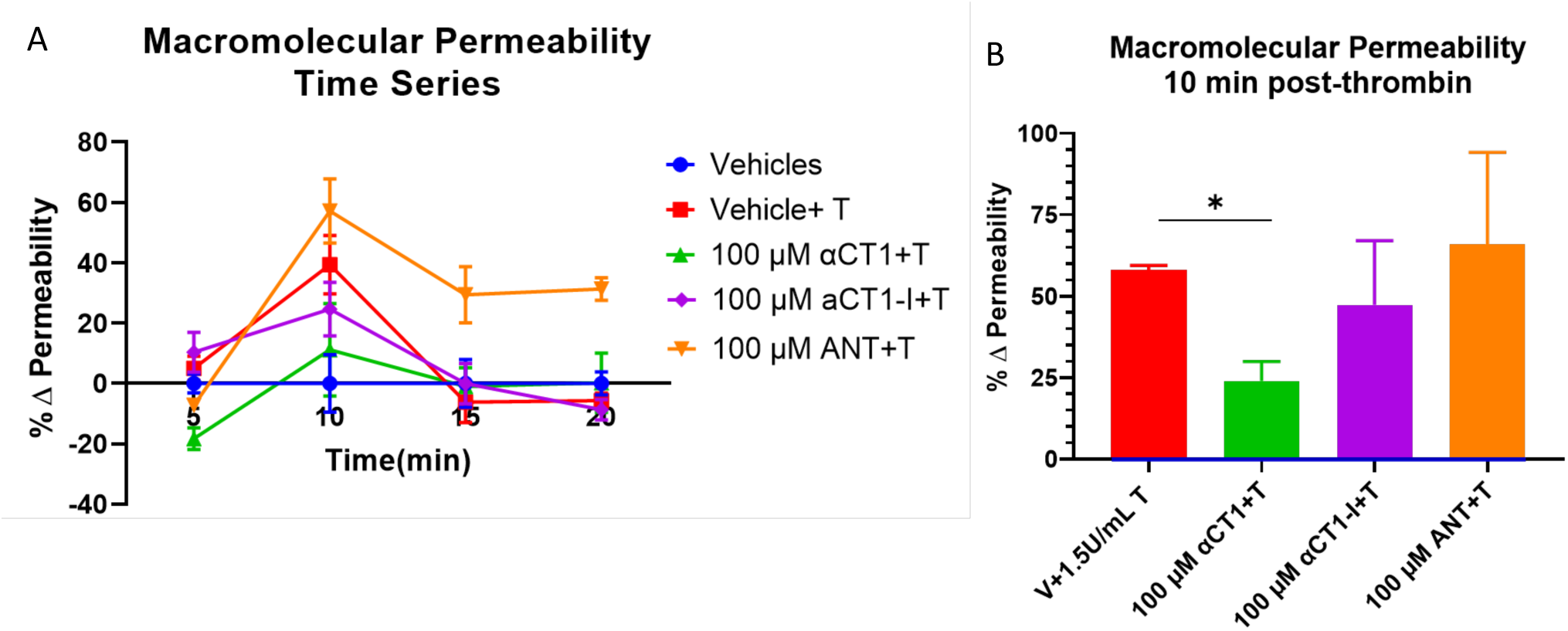
αCT1 requires ZO1 binding-competency to prevent thrombin-induced hyperpermeability in HMEC-1 cell monolayers. **A)** Representative time course of macromolecular flux to 4.5kDa FITC-dextran across the endothelial monolayer, from the top (apical) to the bottom (basolateral) compartment of transwell chambers. The percentage change in absolute permeability was calculated from fluorescent readings of samples taken from the bottom compartment at 5, 10, 15, and 20 min post-thrombin stimulation. Measurements at each time point were normalized to vehicle control. **B)** The change in permeability at the time point of maximum thrombin disruption (10min), normalized to vehicle control, were averaged across experiments. P < 0.05 vs. Vehicle control; N = 4.

### αCT1 prevents thrombin-induced changes in endothelial F-actin and VE-cadherin distribution in a ZO1 binding-competent manner

The mode-of-action of thrombin in disrupting barrier function is thought, in large part, to occur via its effects on the actin cytoskeleton and VE-cadherin (Aslam et al., 2014; Breslin, Zhang, Worthylake, & Souza-Smith, 2015). Furthermore, a previous collaborative report demonstrated that αCT1 produces significant changes to the actin cytoskeleton in brain endothelial cells, via a ZO1 PDZ2 interaction (Chen et al., 2015). Therefore, HMEC-1 cells grown on solid substrates were immunolabeled for F-actin and VE-Cadherin following a similar treatment protocol described for ECIS barrier function experiments, then fluorescent signals imaged using confocal microscopy.

We used this approach, together with a high-throughput quantitative image analysis software, Cell Profiler (McQuin et al., 2018), to quantify changes in the cellular distribution of F-actin and VE-Cadherin in our thrombin/peptide treatment model. Initial observations showed that untreated control HMEC-1 cells grown in monolayers exhibited thin, well-delineated bands of cortical actin marking the boundaries of cells, consistent with intact barrier function, as well as isolated fibers stretching across the cytoplasm of the cell (Figure 5). Thrombin treatment attenuated this sharp F-actin border, causing cells to form densely packed fibrous sheets of stress fibers that stretched across the cell, either through the center of the cell or just outside the cell-center, in a manner consistent with cytoskeletal structures commonly linked to endothelial barrier function disruption in the literature (e.g., Figure 5). Thrombin also increased the formation of intercellular gaps. The thrombin-induced effects on F-actin at 5 min post-thrombin stimulation observed here are consistent with previous reports (Doggett & Breslin, 2011; Rabiet et al., 1996). As for VE-Cadherin, thrombin attenuated the sharp, linear VE-Cadherin signal at the cell border, while simultaneously reducing concentrations of signal towards the center of the cell (Figure 5).

**Figure 5:**
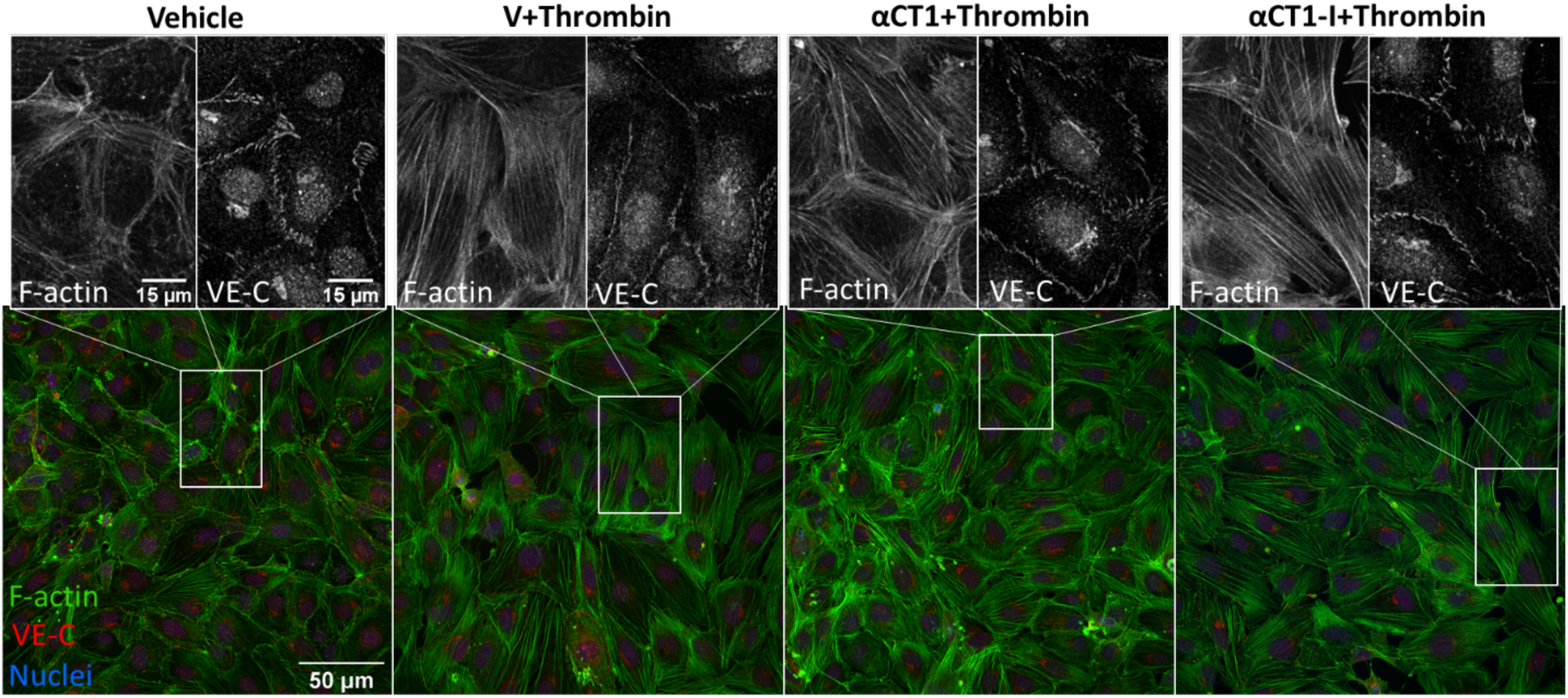
αCT1 inhibits thrombin-induced shift in endothelial F-actin and VE-cadherin distribution in HMEC-1 cell monolayers. Representative confocal images of F-actin and VE-Cadherin in 100µM labeling in peptide-treated, HMEC-1 cells, fixed 5 min after thrombin addition to the media. Zoomed sections show representative treatment-induced F-actin and VE-Cadherin changes.

To quantify these changes, normalized intensities of F-actin and VE-Cadherin immunolabeling were measured at 20 successive equivalently spaced intervals from the nucleus to the peripheral border of cells in the different treatment groups, as detailed in methods (see diagram in Figure 6A). Statistically significant differences in the cellular distribution of F-actin and VE-Cadherin between the treatment conditions compared to thrombin alone are displayed in Figure 6 and in Supplemental Table 1. Overall, F-actin distribution increased more or less linearly from the cell nucleus outward to the cell periphery, peaking in mean fractional intensity near the cell periphery (Figure 6A). VE-Cadherin distribution showed an opposite trend, though with an upward inflection in fractional intensity in region 17, located just a few radii inward from cell borders. Importantly, αCT1, but not αCT1-I pretreatment inhibited the thrombin-induced changes in F-actin morphology, consistent with barrier function effects described previously (Figure 6B). This αCT1-associated effect, compared to αCT1-I, was marked by a significant increase in the proportion of peripherally located cortical actin, simultaneous with increase in VE-Cadherin at cell-cell borders. See Figures 6A and 6B for mean fractional intensity values, and Figures 6C and 6D for vehicle baseline-subtracted values for F-actin and VE-cadherin respectively. αCT1, but not αCT1-I, also prevented thrombin-induced reduction in VE-Cadherin cellular distribution in the four peripheral-most cell compartments, while changes were not significant around the cell center – as indicated by the yellow-highlighted regions on the graphs shown in Figure 6. These data indicated that αCT1 required ZO1-binding competency to protect against thrombin-induced barrier function-associated changes in F-actin and VE-Cadherin.

**Figure 6:**
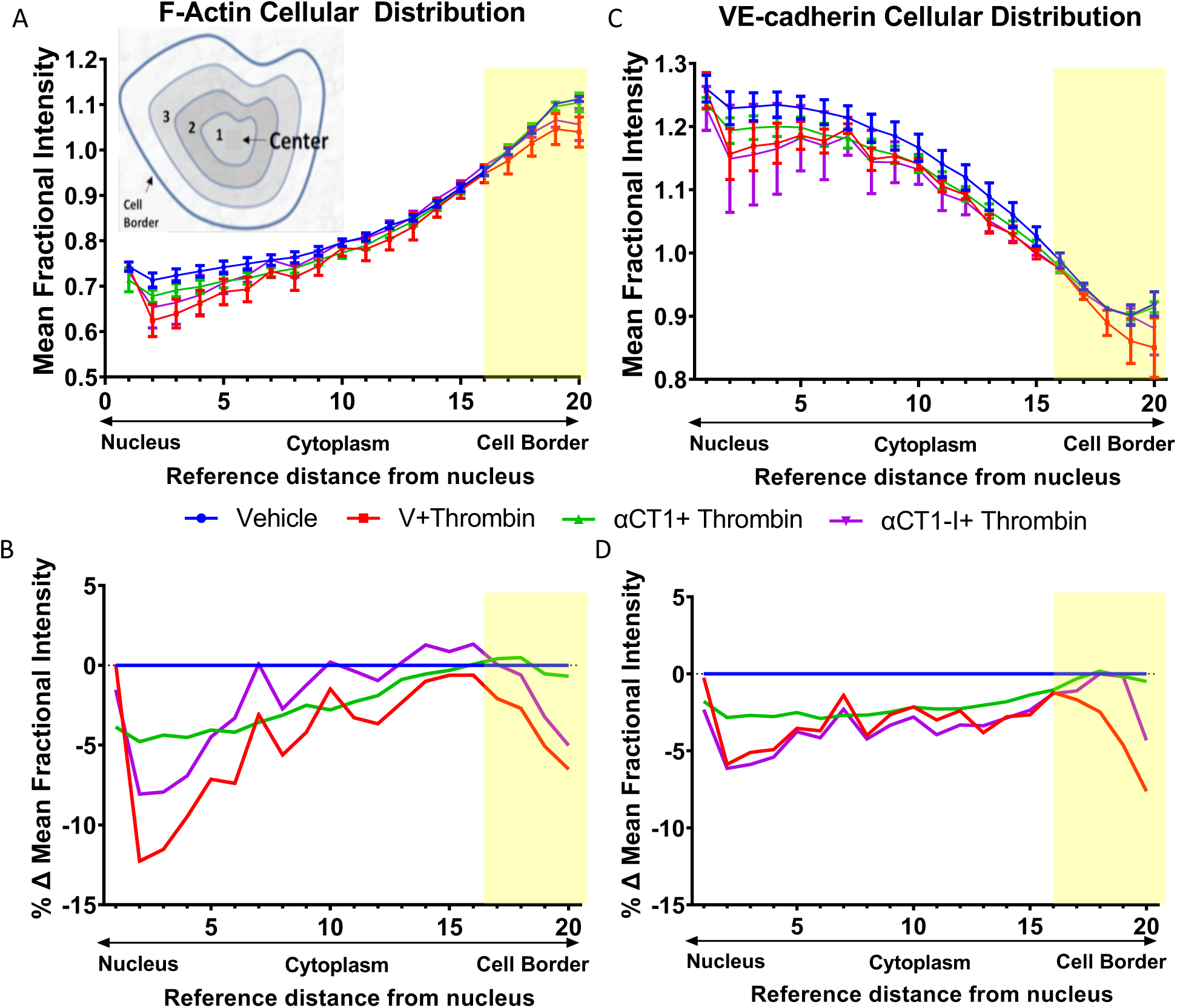
Quantification of αCT1 inhibition of thrombin-induced shift in endothelial F-actin and VE-cadherin distribution in HMEC-1 cells. **A**,**C)** The radial distributions of F-actin (Top left) and VE-cadherin (Top right) were measured as the mean fractional intensity at a given cell radius, calculated as fraction of total intensity normalized by fraction of pixels at a given radius. Cell diagram of regions 1-20 indicated in top left figure. **B**,**D)** F-actin (Bottom left) and VE-cadherin (Bottom right), vehicle-subtracted values calculated as percentage difference from vehicle 100% (Value-Vehicle)/Vehicle). N = 3 Yellow highlighted bar indicates where αCT1+T, but not αCT1-I+T is significant compared to thrombin alone.

### αCT1 requires ZO1 binding-competency to modulate the distribution of F-actin, ZO1 and Cx43 in cultured endothelial cells

To further validate our observations on HMEC-1 cells, another endothelial cell line, Human Dermal Microvascular Endothelial cells (HDMECs), was grown to confluence on collagen-coated transwell filters. HDMECs were used for purposes of improved imaging and quantification of peptide treatment-associated phenomena due to the well-defined cell-cell borders and more uniformly arrayed junctional structures found in this endothelial cell line. Previous reports have indicated that αCT1 targets TJ protein, ZO1, to increase gap junctional Cx43 levels at the cell border in Hela cells, and that gap junctional Cx43 provides points of close cell-cell contact (Elias, Wang, & Kriegstein, 2007; Rhett, Jourdan, & Gourdie, 2011). Therefore, in this set of experiments, cells were stained for F-actin, ZO1, and Cx43 (Figure 6). As in HMEC-1s, untreated vehicle control HDMECs exhibited thin, clearly delineated bands of cortical F-actin marking the boundaries of the cell, while thrombin treatment attenuated this sharp F-actin border, inducing the formation of densely packed stress fibers stretching across the cell, including the cell center. Also similar to the pattern observed in HMEC-1 cells, αCT1, but much less so αCT1-I, blocked this shift in F-actin distribution in HDMECs (Figures 7, 8A-B Supplemental Table 2), with a marked attenuation and enhancement of cytosolic and peripheral F-actin distribution, respectively. HDMEC monolayers pretreated with the cell penetration sequence control, antennapedia (ANT), showed a near identical F-actin distribution pattern as thrombin treatment alone (Figure 8A-B).

**Figure 7:**
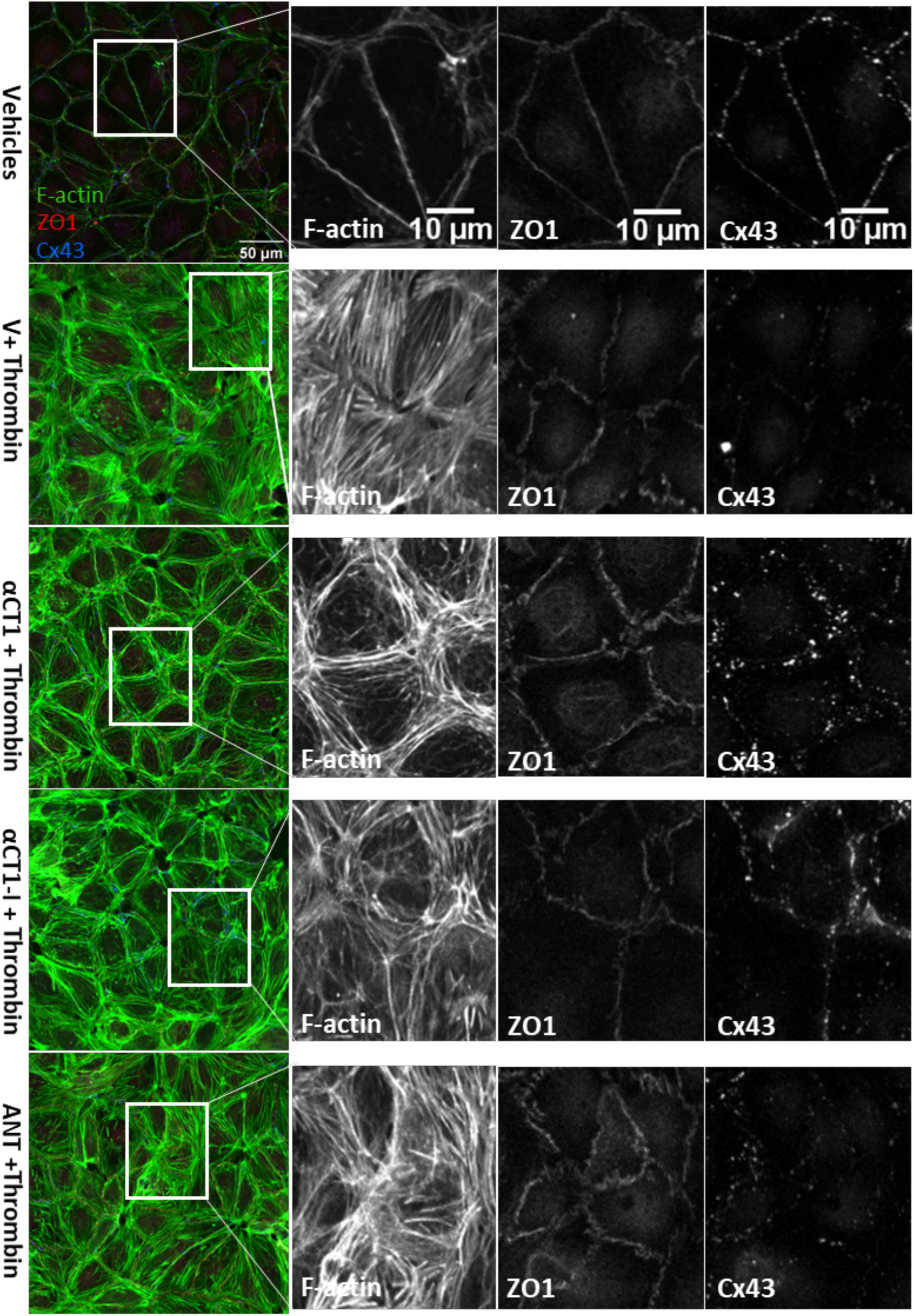
αCT1 inhibits thrombin-induced shift in endothelial F-actin distribution in association with ZO1 and Cx43 remodeling in HDMEC monolayers. **–** Representative confocal images of F-actin cytoskeleton, Cx43, and ZO1 distribution in 100 µM peptide-treated, transwell filter-grown HDMECs, fixed 10 min after thrombin addition to the media. Black and white zoomed images show treatment-induced changes in more detail at the level of cell-cell contacts.

**Figure 8:**
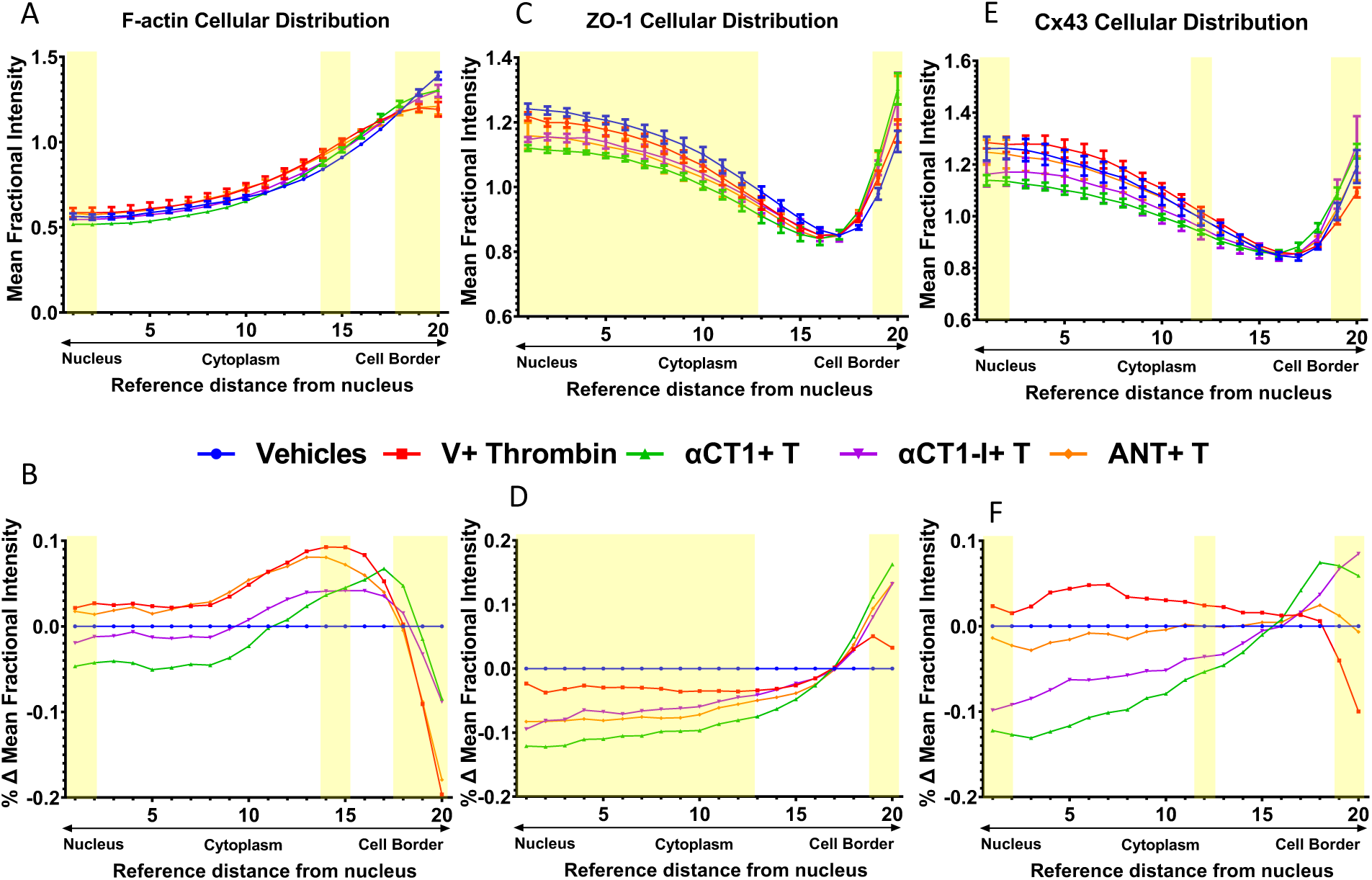
Quantification of αCT1 inhibition of thrombin-induced shift in endothelial F-actin distribution in association with ZO1 and Cx43 remodeling in HDMECs. **A, C, F)** The radial distributions of F-actin (Top left) and ZO1 (Top middle) and Cx43 (Top right)) were measured as the mean fractional intensity at a given cell radius, calculated as fraction of total intensity normalized by fraction of pixels at a given radius. Cell diagram of regions 1-20 indicated in top left figure. **B, D, E)** F-actin (Bottom left) and ZO1 (Bottom middle) and Cx43 (Bottom right), vehicle-subtracted values calculated as percentage difference from vehicle 100% (Value-Vehicle)/Vehicle). N=3. Yellow highlighted bar indicates where αCT1+T, but not αCT1-I+T or ANT+T is significant compared to thrombin alone.

Changes in the distribution of Cx43 and ZO1 induced by thrombin alone did not reach statistical significance at any sub-region within HDMECs (Figures 8C-F, Supplemental Table 2). However, the effects of αCT1 on Cx43 and ZO1 in combination with thrombin were significant, with marked discrimination from the effects of the ZO1 binding-incompetent control, αCT1-I, and the cell penetration sequence control, ANT. Consistent with previous reports on peptide-induced changes in Cx43 distribution at cell-cell contacts (Rhett et al., 2011), αCT1, but not αCT1-I, produced a significant increase in the proportion of Cx43 at cell-cell borders, while both peptides reduced the proportion of signal located in nuclear and cytoplasmic regions (Figure 8B and 8E, Supplemental Table 2). Similar changes in ZO1 signal across the different cellular sub-regions were seen with αCT1, but not αCT1-I or ANT (Figure 8C and 8D, Supplemental Table 2).The 95% confidence intervals for each treatment mean at each sub-region for actin, ZO1, and Cx43 and details of statistically significant differences for these proteins between the treatment conditions compared to thrombin alone are summarized in Figure 8 and Supplemental Table 2. The yellow bars in Figure 8 indicate cell regions in which effects of αCT1 on the junctional protein distributions are discriminated from αCT1-I with respect to thrombin alone (p<0.05). In sum, it was observed that the ZO1-binding-competent Cx43 CT peptide inhibited induction of stress fibers and junctional remodeling in response to thrombin, maintaining actin in more homeostatic-like cortical distributions in the two endothelial cell lines studied.

### αCT1 requires ZO1 binding-competency to exert changes in distribution of cell orientations

An F-actin cytoskeleton-related phenomenon that has been recently linked to barrier function regulation is cellular orientation or handedness (Fan et al., 2018). We assessed cellular orientation on HDMEC monolayers treated with αCT1, as compared to thrombin and peptide controls, and noted that distribution of cell orientations showed significant correlation to the different patterns of actin remodeling seen in our experimental model (Figure 9). Skewness measurements of cell orientation indicated that thrombin shifts the distribution of cell orientation from one side of a normal distribution to the other. That is, under vehicle conditions, the majority of cell orientations took on “negative” angles with respect to an arbitrary X=0° reference axis, while thrombin stimulation “flipped” the cells to take on positive angles. αCT1 pronouncedly reduced the skewness measure to near zero, indicating a near complete attenuation of cell-orientation bias. A Kolmogorov–Smirnov (KS) test on cell orientation data was performed to further confirm the significance of these findings (data not shown).

**Figure 9:**
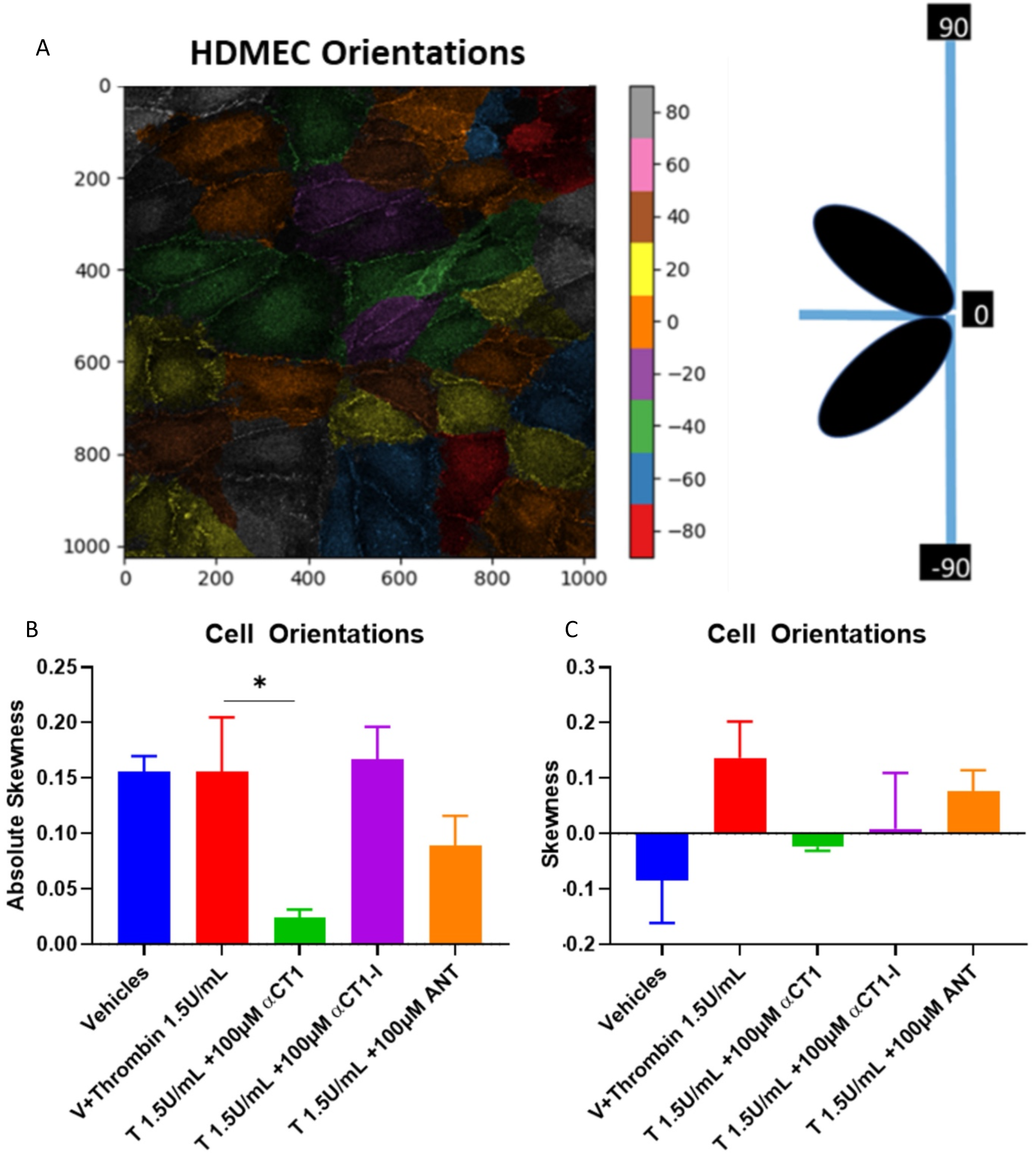
αCT1 alters distribution of cell orientations in HDMEC monolayers. **A)** Representative diagram of cell angle designation across a HDMEC monolayer. **B)** Absolute Skewness measurements, calculated as the absolute value of g1= the average value of z3, where z is the familiar z-score, 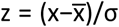, where x is the individual cell angle with respect to 0° angle reference axis. C) Raw Skewness values calculated as g1 above, no absolute value calculated, * P < 0.05 vs. Thrombin; N = 3.

### αCT1 reduces vascular leak in Langendorff-perfused mouse hearts

As mentioned earlier, a cardiovascular protective effect of αCT1 was hypothesized to in part to result from the peptide’s targeting to the coronary vasculature within the heart (Jiang et al., 2019). Thus, to determine if the *in vitro* endothelial barrier protection by αCT1 applied to an *ex-vivo* setting, vascular leakage within peptide-treated mouse hearts was assessed. Langendorff-perfused mouse hearts were perfused for 20 minutes with Tyrode’s solution with or without αCT1 (100uM), followed by 40 minutes with thrombin (1.5U/ml). The permeability tracer FITC-dextran (10 mg/ml), the same tracer used previously in the transwell permeability assay (Figure 3), was added to the final 10 ml of perfusate. Overall, as assessed by quantitative confocal microscopy of cryosections from the hearts, thrombin significantly increased FITC extravasation relative to control (by ∼88%). αCT1 treatment markedly decreased FITC extravasation compared to thrombin alone (p < 0.05 vs. thrombin), nearly restoring it to vehicle control levels (Figure 10).

**Figure 10:**
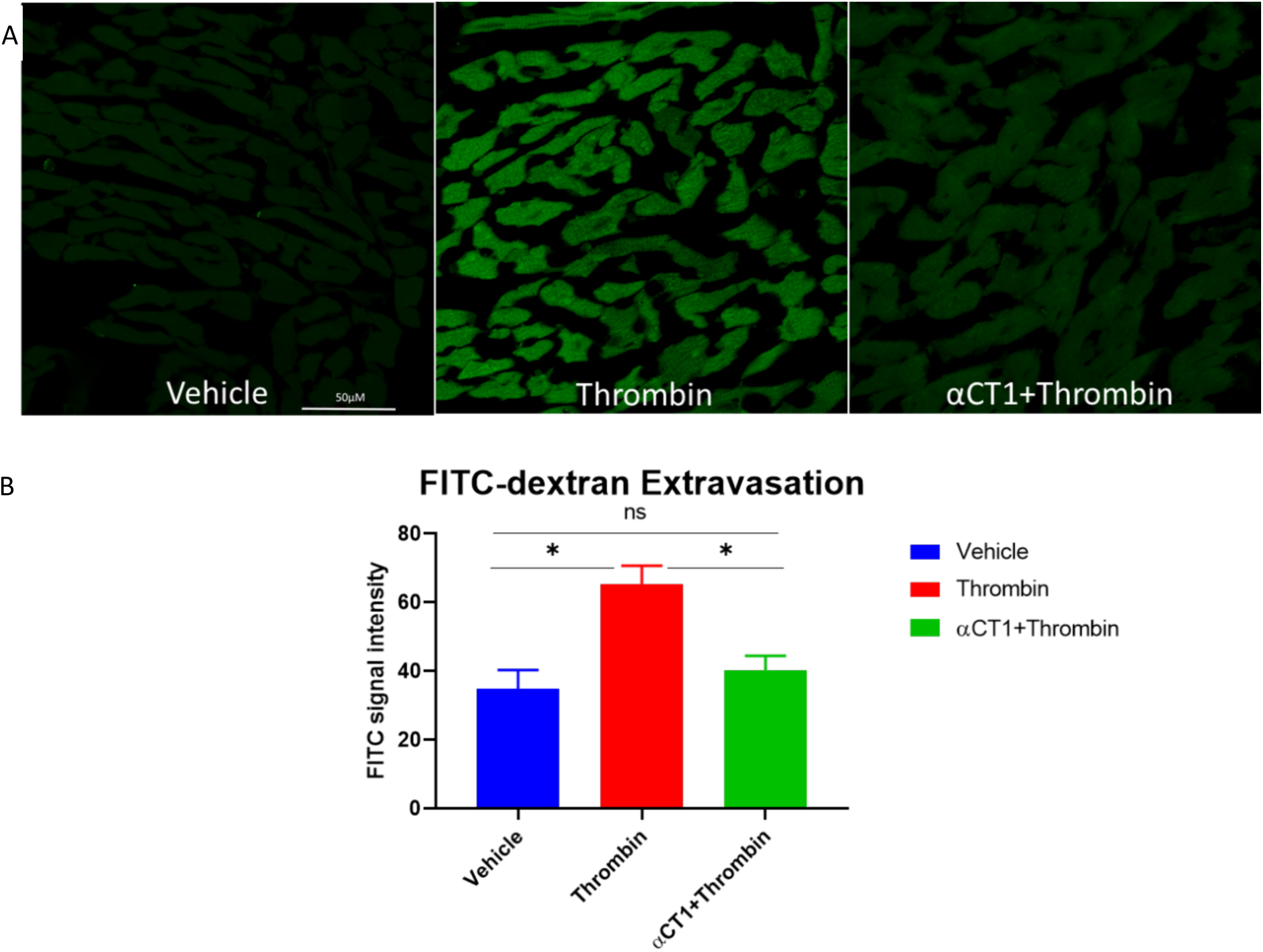
αCT1 inhibits thrombin-intravascular leak in Langendorff-perfused mouse hearts. **A)** Representative confocal images of FITC-dextran extravasation within Langendorff-perfused mouse hearts. **B)** Quantification of FITC-dextran signal within mouse hearts perfused 40 minutes with thrombin (1.5U/ml) with or without 20 min αCT1 (100uM) pre-treatment. SE bars, * P < 0.05

## Discussion

In this study, we investigated the effects of Cx43 CT mimetic peptide αCT1 on trans-endothelial permeability and junctional and cytoskeletal proteins that determine this function. We found that pre-treatment with αCT1 protects vascular barrier function from thrombin-induced disruption in *ex vivo* (Langendorff-perfused mouse heart) and *in vitro* (ECIS and Transwell permeability) models. Barrier protection *in vitro* by αCT1 occurred in association with localization of the peptide with ZO1 at cell-to-cell borders, specific effects on cell orientation and changes in patterns of F-actin, VE-Cadherin, Cx43, and ZO1 remodeling, particularly at the periphery of cells. Importantly, a ZO1 binding-incompetent variant of αCT1, αCT1-I, showed no propensity to associate with ZO1 at the cell periphery and also demonstrated no facility for protecting barrier function, suggesting that ZO1 binding-competency is required for Cx43 CT mimetic peptides to affect the vascular permeability parameters assessed.

The findings we present herein are consistent with previous studies indicating a protective role of Cx43 CT in channel-independent modulation of barrier function. Mice deficient in the Cx43 CT die as a result of epithelial barrier dysfunction, despite maintaining normal GJIC (Maass et al., 2004). Obert and colleagues (2017) showed that the Cx43 CT mimetic, αCT1 prevented breakdown of TJ-based barrier function via a channel-independent mechanism, in Cx43-expressing epithelial cell lines derived from the retinal pigment layer (Obert et al., 2017). Three novel insights from the present study are that: 1) In addition to protecting epithelial cell barriers, αCT1 is protective of barrier function in endothelial cells; 2) The terminal isoleucine of αCT1, and thus maintenance of the peptide’s high affinity interaction with the PDZ2 domain of ZO1, appears to be required for barrier protective properties in the models studied; and 3) The inhibitory effect of αCT1 on actin remodeling in response to a stressor such as thrombin appears to be central to the activity of the peptide in barrier function protection.

A major finding of this study is that αCT1 inhibits thrombin-induced attenuation of cortical actin and F-actin stress fiber formation. In intact endothelial barriers, cortical actin, in association with junctional complexes, exert outward directed tension between cells, in dynamic balance with opposing inward contractile forces within cells. The actin stress fiber phenotype induced by thrombin shifts the balance of forces within and between cells resulting in a disruption of cell contacts, formation of extracellular gaps and breakdown of barrier properties (Aslam et al., 2014; Belvitch et al., 2018; Chugh & Paluch, 2018; Escribano et al., 2019; Shakhov, Dugina, & Alieva, 2019 – see also Figure 11). In addition to thrombin, numerous other chemical and physical stressors, including histamine, lipopolysaccharide, endotoxin, Tissue Necrosis Factor (TNF), and shear stress, cause similar shifts in the balance of intra- and intercellular forces, together with loss of barrier patency via the same actinomyosin-based mechanism. For example, Mehta and colleagues (2002) showed that pretreating Human Pulmonary Artery Endothelial Cells (HPAEC) with latrunculin-A (Lat-A), a toxin known to prevent F-actin assembly, inhibited thrombin-induced endothelial cell retraction and decreased loss of transepithelial electrical resistance (TEER) (Mehta et al., 2002). Pertinent to the current study, a report by Chen and colleagues (2015) found that αCT1 produced derangement of cytoskeletal fibers when applied to brain endothelial cells, including formation of cytoplasmic actin-rich node-like structures (Chen et al., 2015). Interestingly, the authors also reported that these results could be recapitulated by over-expressing a PDZ2 domain-deleted ZO1 mutant, suggesting that the Cx43-binding domain of ZO1 targeted by αCT1 was necessary for the observed effects on actin cytoskeleton organization.

**Figure 11:**
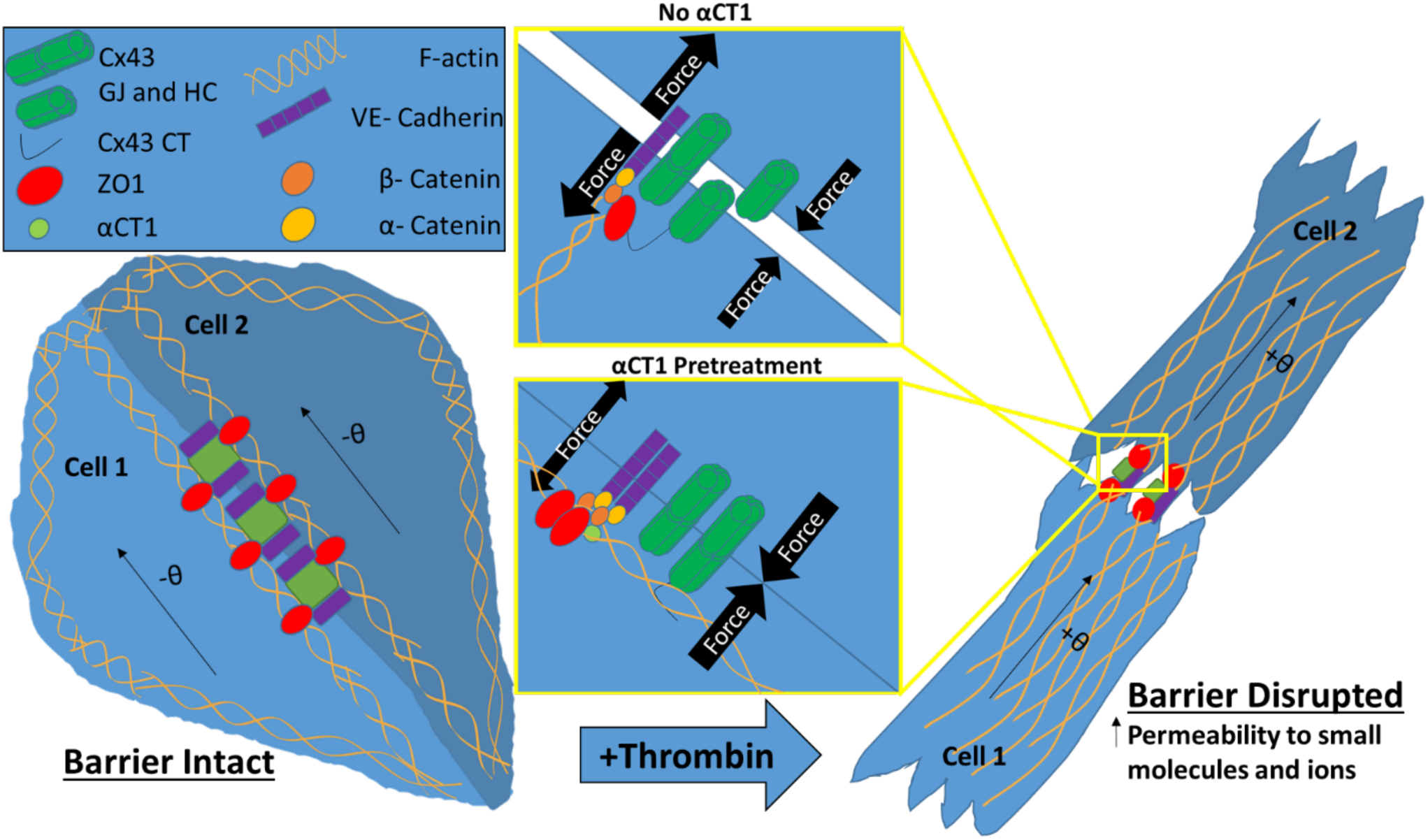
Simplified model of αCT1 effects on endothelial cells in monolayers under thrombin stimulation. Thrombin stimulation of endothelial cells produces a redistribution of F-actin from peripherally located cortical actin to stress fibers that cut across the cytoplasm. These stress fibers terminate onto VE-cadherin containing adherens junctions at the cytoplasmic plaque of the membrane, and in their contractile state, and are thought to exert barrier-destabilizing pulling forces at the cell membranes of opposing cells (Burridge & Wittchen, 2013). This accompanies a reduction and remodeling of adherens junctions, intercellular gap formation and increased permeability to small molecules and ions. In correlation with these changes, thrombin flips the F-actin controlled orientation of cells in the monolayer to take on angles opposite to those observed in homeostatic conditions in which the barrier is intact (Fan et al., 2018). αCT1 pretreatment of endothelial monolayers appears to inhibit these thrombin-induced changes in a ZO1-interaction-dependent manner, reducing stress fiber formation and maintaining actin in more cortical distributions. αCT1 treatment is also associated with increased Cx43 gap junctional contacts, maintenance of VE-cadherin-containing adherens junctions and ZO1-containing tight junctions at cell borders. We hypothesize that uncoupling of ZO1 from anchorage at key membrane-associated partner proteins (e.g., Cx43, α-catenin) via ligand binding to its PDZ2 domain (e.g., by αCT1), may offset the perpendicular alignment of F-actin fibers. This in turn may reduce the ability of stress fibers aligned in this manner to exert centripetal force onto the cytoplasmic face of adherens junctions, reducing extracellular gap formation, stabilizing the endothelial barrier and maintaining heterogeneous patterns of cell orientation in response to stressors such as thrombin.

We have previously demonstrated that pre-treatment with either αCT1 or αCT1-I can reduce the severity of myocardial ischemia-reperfusion (IR) injury (Jiang et al., 2019). These results in myocardium stand in contrast to the apparent mechanism of the selective vascular endothelial barrier protective effect of αCT1 characterized herein. The shared myocardial protective effect αCT1 and αCT1-I occurs independent of ZO1 interaction, and is correlated with negatively charged sequences common to both peptides, which mediate binding to the H2 domain of Cx43 (Jiang et al., 2019). The severity of heart IR injuries is thought in part to be determined by levels of activation of myocardial Cx43 hemichannels (Marsh, Williams, Pridham, & Gourdie, 2021; Schulz et al., 2015) and we have previously proposed that reductions in channel activity associated with targeting the Cx43 H2 domain could account for the cardioprotective effects elicited by αCT1 and αCT1-I. (Jiang et al., 2019). By contrast, increased trans-epithelial permeability in endothelial monolayers subject to a thrombin insult, as studied herein, seems to be primarily mediated via effects on actin organization and shifts in forces exerted on intercellular contacts downstream of this remodeling of the cytoskeleton (Aslam et al., 2014; Vouret-Craviari, Boquet, Pouysségur, & Van Obberghen-Schilling, 1998) Actin’s propensity to interact with ZO1, or membrane bound actin-binding ZO1 partners such as cytoplasmic components of adheren junctions (e.g. α-catenin (Maiers et al., 2013)), and its capacity to form and align stress fibers, appears to be sensitive to a αCT1-induced modulation of ZO1 following exposure of cells to thrombin. That αCT1 treatment resulted in altered patterns of actin cytoskeleton remodeling, and in particular to that of cortical actin at the cell periphery, is also consistent with thrombin’s well-established mode-of-action on vascular permeability (Bogatcheva, Garcia, & Verin, 2002).

Our results indicate that αCT1 inhibits a thrombin-induced reversal of cell orientation, pronouncedly attenuating cell orientation bias in a ZO1 interaction-associated manner, while enhancing ZO1 localization at cell boundaries. Cell orientation distribution has been linked to actin-mediated ZO1-associated barrier integrity in a pioneering study carried out by Fan and colleagues (Fan et al., 2018). These authors determined that endothelial barrier disruption triggered by a PKC activator IndoV, correlated to reduced ZO1 expression and actin-dependent reversal of cell orientation. Skewness measurements of cell orientation undertaken in our study indicate that thrombin shifts the distribution of cell orientation from one side of a normal distribution to the other. While the analysis in the present study was not carried using a well-defined reference axis based on the tangential direction of a micro-patterned circular array as in the study by Fan and co-workers (Fan et al., 2018), our skewness results are consistent with their observations. Ongoing studies may usefully focus on if and how Cx43 and Cx43/ZO1 interactions may operate in this context, potentially contributing to the handedness of actin cytoskeletal and cell orientation responses.

In the current study, αCT1 maintained F-actin at the cell periphery under thrombin stimulation, while at the same time augmenting the border localization of Cx43, ZO1, and VE-Cadherin. It is well established that stabilization of barrier function is often marked by restoration of AJ and TJ proteins to cell-cell borders (Radeva & Waschke, 2018; Riesen, Rothen-Rutishauser, & Wunderli-Allenspach, 2002). Furthermore, multiple studies have demonstrated cell-cell adhesive roles for Cx43 GJ (Cotrina et al., 2008; Elias et al., 2007; Lin et al., 2002), and the upregulation of GJ Cx43 has been shown to promote a stabilization of cortical actin (Francis et al., 2011; Kameritsch et al., 2015; Xu et al., 2006). Under normal conditions, cortical actin promotes the stability of cell-cell interactions by tethering these junctional structures (E.g. GJ, TJ, AJ) with other intracellular components (García-Ponce, Citalán-Madrid, Velázquez-Avila, Vargas-Robles, & Schnoor, 2015; Rodgers, Beam, Anderson, & Fanning, 2013). Taken together, our data suggests that αCT1 protects barrier function first and foremost, by inhibiting a shift in F-actin away from cell-to-cell contacts, thereby stabilizing transcellular interacting proteins, VE-Cadherin and Cx43, and the TJ-scaffolding protein ZO1. Figure 11 provides a model of how αCT1 pretreatment could enhance outward directed tension and minimize inward directed pulling forces via modulation of actin and junctional protein distribution, with downstream effects on endothelial gap formation and barrier permeability.

Further insight into αCT1’s mechanism can be gained from the literature on the role of sphingolipid, Sphingosine-1-phosphate (S1P) in barrier modulation. A report by Want et al using atomic force microscopy showed that thrombin caused a decrease in cortical actin, concomitant with a drop in cell stiffness at the cell border, while S1P had opposite effects (Wang et al., 2015). Moreover, Lee and colleagues (Lee et al., 2006) demonstrated that in association with barrier function stabilization as measured by ECIS, S1P stimulation caused a redistribution of ZO1 and Claudin-5 to cell-cell contacts, and enhanced border colocalizations of ZO1/ cortactin and ZO1/α-catenin in Human Umbilical Vein Endothelial Cells(HUVEC). While no known direct interaction between S1P and ZO1 has been identified to date, we speculate that the CT of Cx43 (either endogenous or exogenously applied in the form of αCT1) and S1P, may share a similar mode-of-action in modulating ZO1/actin-mediated effects on endothelial barrier function.

In addition to utility of αCT1 as a tool for addressing basic research questions about the potential role of Cx43 CT in barrier function, the therapeutic potential of this peptide in the treatment of vascular edema could be considerable. αCT1 has undergone clinical testing in humans for a number of skin-related disease indications, including in healing of chronic wounds, where swelling and edema, and thus disrupted barrier function, are well characterized aspects of pathology (Ghatnekar, Grek, Armstrong, Desai, & Gourdie, 2015; Grek et al., 2017; Grek et al., 2015; Montgomery, Ghatnekar, Grek, Moyer, & Gourdie, 2018). In the present study, αCT1 pronouncedly attenuated vascular leak in Langendorff-perfused mouse hearts. For a large set of disorders (e.g. sepsis, ischemia-reperfusion (IR) injury, major trauma, organ transplantation) and tissue types, organ dysfunction and patient outcomes associates with microvascular dysfunction and edema (Chistiakov, Orekhov, & Bobryshev, 2015; G. Heusch, 2016). In other studies, we have also linked edema, such as that occurs following injury to the heart, to increased propensity to develop deadly arrhythmias (Veeraraghavan et al., 2015; 2018). Given the findings from the present study, αCT1 might be considered a potential vascular-targeting, anti-edema treatment strategy for cardiovascular injury and other diseases in which edematous accumulation is detrimental. Thus, future work might investigate whether or not the ability of αCT1 to inhibit of vascular leakage within the heart, extends to vascular barrier protection in other tissues/organs.

## Materials and Methods

### Test Reagents

Peptides Biotin-αCT1 (Biotin-RQPKIWFPNRRKPWKK-RPRPDDLEI), Biotin-αCT1-I (Biotin-RQPKIWFPNRRKPWKK RPRPDDLE), and Biotin-ANT (Biotin-RQPKIWFPNRRKPWK), were synthesized and quality checked for fidelity and purity using high-performance liquid chromatography and mass spectrometry (LifeTein, Hillsborough, NJ). Thrombin was purchased from Millipore Sigma (Burlington, MA Cat: T7513)

#### FITC-dextran extravasation

Langendorff-perfused mouse hearts were perfused for 20 minutes with Tyrode’s solution with or without αCT1 (100uM), followed by 40 minutes with thrombin (1.5U/ml). FITC-dextran (10 mg/ml) was added to the final 10 ml of perfusate. Perfused hearts were then cryopreserved as described above and extravasated FITC-dextran levels assessed by confocal microscopy of cryosections.

### Impedance measurement using ECIS

The barrier integrity of HMEC-1 (CDC, Atlanta, GA) was measuring using ECIS Z Theta system (Applied Biophysics, Troy, NY). HMEC-1 monolayers with a seeding density of 7.50×10^4^ cells/cm^2^ were grown to confluence (24-72 h) on collagen I coated 8 W 10E+ electrodes. After cell sedimentation and attachment to the electrode surface within 30 min at room temperature, the 8-well arrays were placed inside the ECIS® device for impedance monitoring. All ECIS® measurements were analyzed at an AC frequency of 32 kHz, which was identified as the most sensitive frequency for this cell type (e.g. frequency at which maximum difference between cell-containing and cell-free measurements was achieved), each well measured every 2-4 minutes. 1h prior to treatment, media was exchanged with 360µL fresh media. Test reagents were diluted in pre-warmed medium. 20µL peptide/media solution (αCT1, αCT1-, ANT) was added to a final concentration of 100µM. Cells incubated in peptide for 1 hr, then 20µL thrombin/FBS-free media solution was added to a final concentration of 0.5U/mL. Approximately 5 min following thrombin addition, peptide-induced barrier function protection was calculated as the percentage of barrier protection from thrombin disruption=[(ohmic resistance peptide-ohmic resistance of thrombin)/ (ohmic resistance thrombin – ohmic resistance vehicle control)) x 100%. The effects of peptides alone on barrier function were calculated as the change in ohmic resistance compared to vehicle control (ohmic resistance peptide-ohmic resistance vehicle).

### Macromolecular permeability (MP)

Macromolecular permeability (MP) filter inserts (pore size 0.4 μm, 12 mm diameter) (Falcon, Corning, NY; Cat:353095) were coated with collagen I (Corning, Corning, NY; Cat: 354246) at 2µg/mL in 0.02N acetic acid. Subsequently, the lower compartments of 24W Transwell chambers (Falcon, Corning, NY; Cat: 353504,) were filled with 700µL HMEC-1 media. HMEC-1 cells suspended in 300 μl media (7.50×10^4^ cells/cm^2^) were seeded on the upper compartment. They were grown to confluence (48–96 h). Cells were treated as indicated in the ECIS experiments (see above). At 55 min after peptide addition, 4µL 100mg/mL FITC-dextran/H_2_0 solution was added to wells. 150microl of basolateral media was collected for time 0. Thrombin was added to experimental wells to final concentration of 1.5U/mL, 5 min after the application of FITC-dextran. 150 μl samples were taken after 5, 10, 15, 20 min from the lower compartment. The removed volume was immediately replaced by fresh medium. To evenly disperse FITC-dextran within the media, transwell plates were gently shaken. Fluorescence (ex: 485 nm; em: 535 nm) was measured with a fluorescence plate reader. Data are expressed as relative changes in fluorescence compared to vehicle permeability. P (cm/s) was calculated by the following equation(Bischoff et al., 2016).

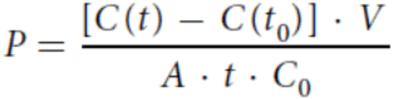

C(t) is the concentration (μg/ml) of FITC-dextran in the samples that were taken from the lower compartment after 5, 10, 15, 20 min, C(t0) is the FITC dextran concentration (μg/ml) of the samples taken after 0 min, t is the duration of the flux (s), V is the volume (cm^3^) in the lower compartment, A is the surface of the Transwell membrane (cm^2^) and C0 is the initial concentration (μg/ml) of the tracer on the dornor side. The concentration of FITC-dextran in each sample was determined by reference to a FITC-dextran standard curve.

### Proximity Ligation Assay

The peptide/ZO1 interaction was detected *in situ* using the Duolink secondary antibodies and detection kit (Sigma, St Louis, MO, Cat: 92002, 92004) according to manufacturer instructions. Primary antibodies against Cx43 (South San Francisco, CA; Cat: SC6560) and biotin (Invitrogen, Carlsbad, CA ; Cat: 617-300) were applied under standard conditions. Duolink secondary antibodies against the primary antibodies were then added. These secondary antibodies were provided as conjugates to oligonucleotides that when within close proximity(< 40nm; ; Gullberg, 2010) were ligated together Duolink Ligation Solution. Finally, polymerase was added, to trigger closed circle rolling amplification(which amplified any existing closed circles) and detection was achieved with complementary, fluorescently labeled oligonucleotides.

Confocal images were acquired on a TCS SP8 laser scanning confocal microscope (LSCM) equipped with a 63×/1.4 numerical aperture (NA) oil objective (Leica, Buffalo Grove, IL). Imaging processing and quantitative image analysis of Duolink signal was done using Cell Profiler (MIT, Cambridge, MA). An intensity threshold was applied, then object clusters between 2 and 50 pixels in diameter were identified as Duolink Objects, then compared to original signal for validation. These Duolink objects were then counted and normalized to the number of nuclei in the images, and these values were normalized again to No Peptide Control.

### Immunostaining and Quantitative Image Analysis

For peptide uptake experiments (Figure 1), Cx43-deficient MDCK monolayers were treated with peptide for 1 hr, then cells were washed with DPBS w/Ca2+ and Mg2+, then fixed with 4% paraformaldehyde. The biotin portion of the peptides were labeled with Streptavidin, Alexa Fluor 647 (Invitrogen, Carlsbad, CA ; Cat: S21374). ZO1 was detected with Rb A-ZO1 (Zymed, South San Francisco, CA; Cat: 61-7300) and Chicken A-Rb 488 (Life Technology, Carlsbad, CA; Cat: A21441). Actin was labeled in HDMEC, grown to confluence on transwell filters (Promocell, Heidelberg, Germany) by Alexa Fluor 647 phalloidin (Invitrogen, Carlsbad, CA; Cat: A22287)(Figure 3). Colocalization analysis was done by isolated border ZO1 pixels and calculation Pearson correlation coefficient with peptide biotin signal. For the distribution of cell orientations in HDMECs, absolute skewness measurements were calculated as the absolute value of g1= the average value of z^3^, where z is the familiar z-score, 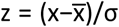, where x is the individual cell angle with respect to a 0° angle reference axis. Quantitative Image Analysis of F-actin, VE-Cadherin, ZO1, and Cx43 was done using Cell Profiler (MIT, Cambridge, MA) (McQuin et al., 2018). For a given cell within a monolayer selected at random to be quantified, a mask based on immunolabeling signals was created in Cell Profiler. Then, the radial distribution of relative F-actin, Cx43, ZO1 or VE-Cadherin labeling levels were measured from the cell center to the cell border in 20 successive sub-regions. The sub-regions were each ∼1-1.5 µm width. Normalized fractional intensity for each sub-region was calculated as a fraction of total intensity normalized by fraction of pixels at a given radius. Means at each cell location were then estimated in R Studio (R Core Team, 2021) by employing a General Linear Model with Random Effects to account for the variability within each cell location. 95% confidence intervals for each treatment mean at each cell location were then calculated.

### Statistics

All data from at least three independent experiments are presented as mean± standard error of the mean (Alonso et al.). Statistical significance was evaluated using GraphPad Prism (version 8.3, GraphPad Prism, San Diego, USA) and assessed by one-way ANOVA and post-hoc tests properly corrected for multiple comparisons where applicable. For protein radial distribution statistics, a General Linear Model with Random Effects was utilized in R Studio, (R Core Team, 2021) and estimated means from the model were calculated with 95% confidence intervals. Significant differences between treatments groups and thrombin treatment alone were reported in Tables 1 and 2. A Kolmogorov–Smirnov (KS) test on cell orientation data was performed to confirm the significance of these findings (data not shown). Significant differences were assumed at *P* ≤ 0.05.

## Acknowledgements

We would like to thank Jane Jourdan for her technical assistance with some of the experiments conducted in this manuscript. Ian Crandell(PhD) and Jennifer West(MS), are thanked for their expert guidance on a few of the statistics components of the manuscript.

## Author Contributions

Conceptualization and design, experimental investigation and writing, R.E.S.; Conceptualization and design, writing—review and editing, R.G.G. Experimental contributions, LM and RV. All authors have read and agreed to the published version of the manuscript.

## Supplementary Figures

**Supplementary Figure 1:**
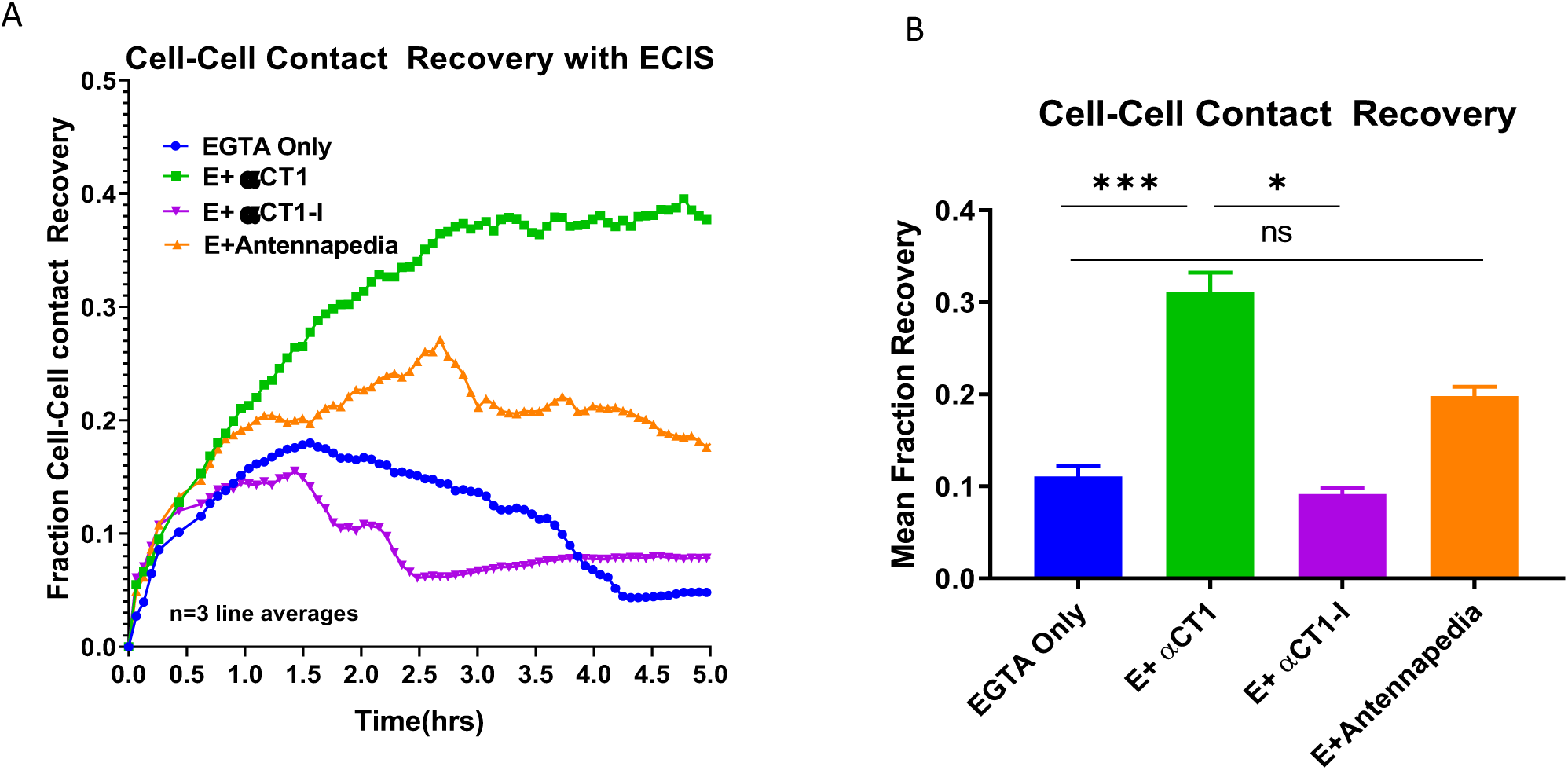
αCT1 augments barrier function recovery in Cx43-deficient MDCK cells. **A)** Summary ECIS time course data of barrier function recovery calculated as fraction of barrier function recovery, normalized to the difference between baseline and time points of maximal disruption. **B)** Quantification of area under the curve analysis of barrier function recovery across 5 hour time period, applied to time course data from Figure 4A.

**Supplementary Table 1:**
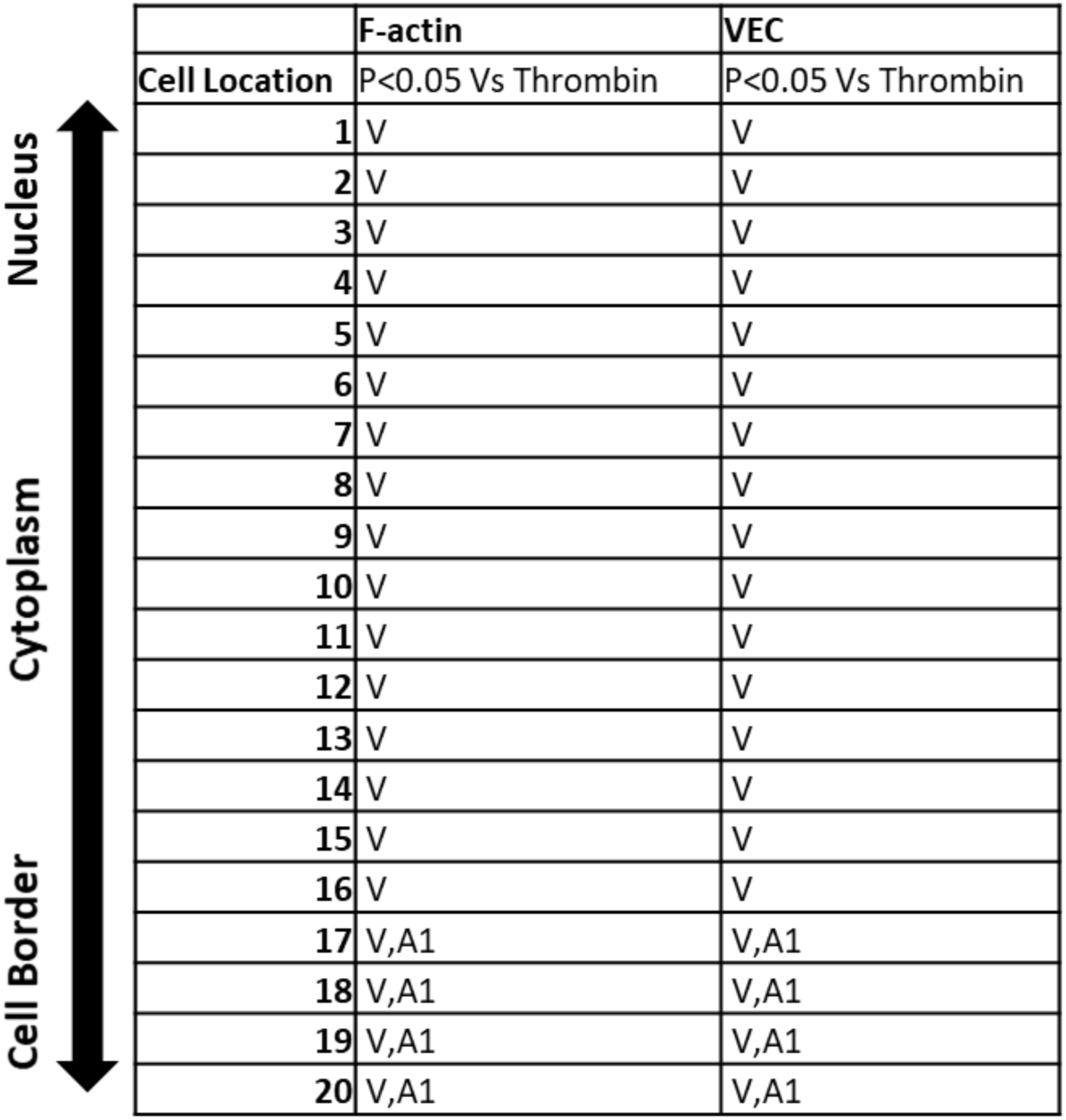
Peptide treatments with significant effects compared to thrombin treatment alone in HMEC-1. **V**=Vehicle **A1**=αCT1+T **A-I**= αCT1-I +T

**Supplementary Table 2:**
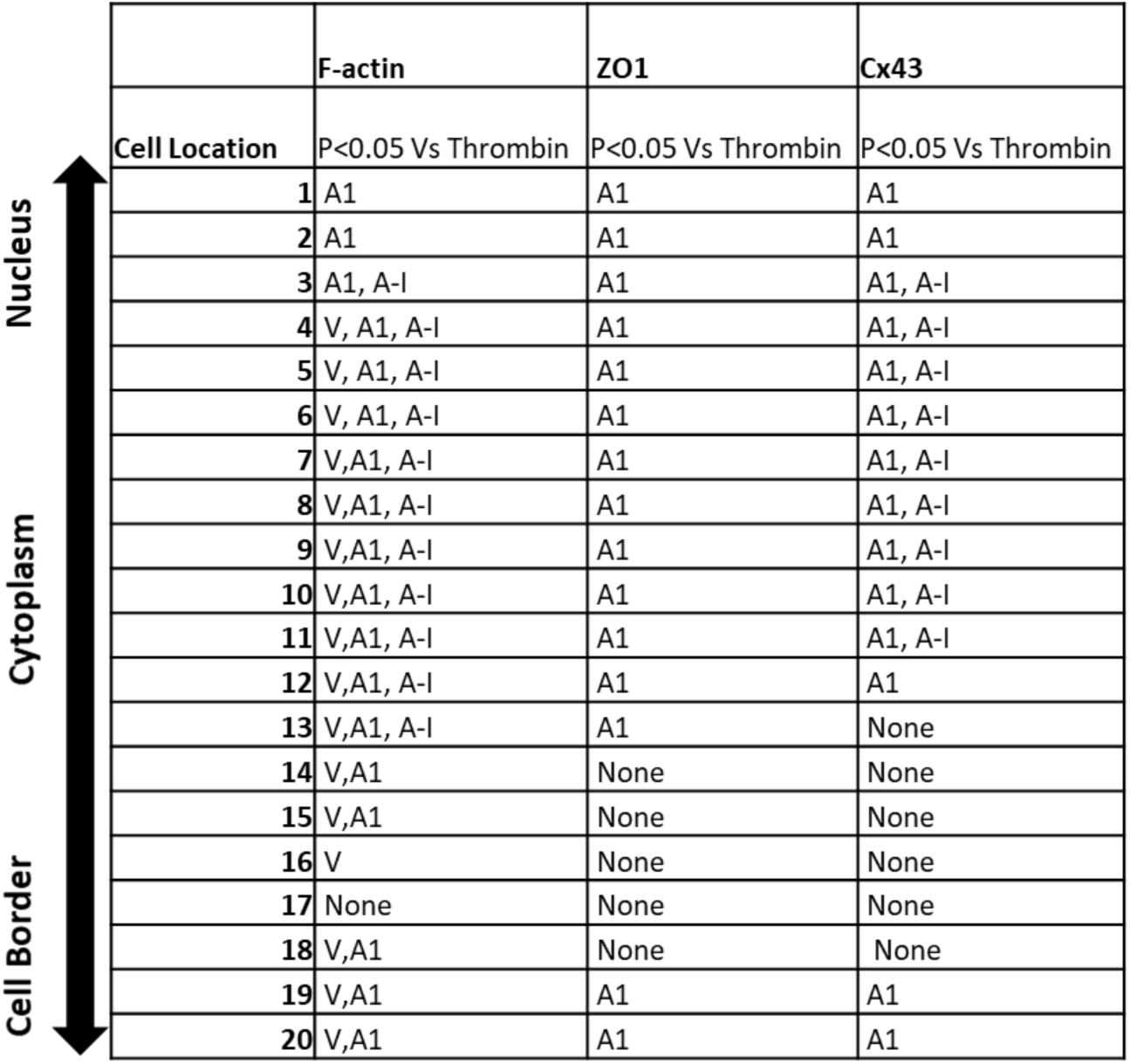
Peptide treatments with significant effects compared to thrombin treatment alone in HDMECs. **V**=Vehicle **A1**=αCT1+T **A-I**= αCT1-I +T **ANT**= ANT +T

